# Protein structure-informed deep learning enables species-specific codon optimization

**DOI:** 10.64898/2026.04.21.720047

**Authors:** Wenzhe Jin, Wuwei Tan, Hui Li, Xiangyu Ji, Ming Li, Deqiang Zhang, Jinbo Xu

## Abstract

Codon usage bias is highly species-specific, posing a major challenge for heterologous protein expression. Existing deep learning approaches to codon optimization rely primarily on DNA or protein sequence information and largely neglect constraints imposed by protein structure and folding. Here, we present Protein structure–Informed Species-specific Codon Optimization (PISCO), a Geometric Vector Perceptron (GVP)–based model that integrates protein sequence, three-dimensional protein structure, and host codon usage statistics to generate optimal, host-specific codon sequences. Compared with protein-structure–agnostic models, PISCO improves codon recovery by 6% and substantially increases similarity to natural coding sequences, reducing divergence by at least 42% in Codon Similarity Index (CSI), 50% in Codon Frequency Distribution (CFD), and 14% in Dynamic Time Warping (DTW) metrics. Ablation analyses demonstrate that incorporating protein folding kinetics and host-specific information is critical to these gains. Moreover, by leveraging host codon usage statistics, PISCO generalizes to optimize codon sequences for species absent from the training data. An autoregressive variant of PISCO further enhances concordance with natural codon usage patterns, at the cost of a modest reduction in codon recovery rate. Wet-lab validation confirms that PISCO-optimized sequences significantly enhance protein solubility and functional expression. Together, these results establish protein structure as a key determinant of species-specific codon optimization and provide a transferable framework for structure-aware gene design.

## 1 Introduction

Efficient heterologous protein expression remains a central challenge in synthetic biology and biotechnology. While the genetic code is redundant, the non-uniform usage of synonymous codons—known as codon usage bias (CUB)—exerts a profound influence on translation efficiency and protein biogenesis [1–4]. Traditionally, codon optimization strategies have focused on maximizing translation speed by replacing rare codons with high-frequency ones, under the assumption that increasing elongation rates directly boosts protein yield. However, growing evidence suggests that this “speed-centric” view is incomplete. Proper folding of nascent polypeptides frequently requires regulated translation dynamics, including programmed ribosomal pausing facilitated by rare codons [5–7]. Consequently, a fundamental tension exists: while fast translation favors high expression, it may simultaneously lead to protein misfolding and aggregation.

Over the past decades, codon optimization strategies have evolved in tandem with an increasingly nuanced understanding of translational mechanisms. Initial approaches relied on simple statistical heuristics based on natural codon frequencies, as exemplified by methods such as Uniform Random Choice (URC), Background Frequency Choice (BFC), Extended Random Choice (ERC), and Highest Frequency Choice (HFC). To better support protein folding, subsequent developments introduced “codon harmonization” [8–11], which seeks to preserve the original translation rhythm of the source species rather than merely maximizing elongation speed. More recently, the field has shifted toward deep neural networks that integrate the entire Coding DNA Sequence (CDS) to capture complex, long-range dependencies across the transcript [12–14]. These frameworks have further evolved to incorporate host-specific information and regulatory contexts, such as 5’ and 3’ untranslated regions (UTRs), to improve crossspecies generalizability [15–18]. While some recent models like TransCodon have begun to consider structural features, such as 5’ UTR secondary structures [18], existing methods remain primarily sequence-centric. They largely fail to explicitly account for the three-dimensional structural constraints of the emerging nascent chain, which fundamentally limits their ability to model co-translational folding, a process inherently governed by structural constraints.

We hypothesize that effective codon design requires a holistic integration of three key factors: (1) translation kinetics encoded by codon usage, (2) co-translational folding signals determined by protein structure, and (3) host-specific constraints. To test this, we propose **P**rotein structure **I**nformed **S**pecies-specific **C**odon **O**ptimization (**PISCO**), a unified framework that explicitly incorporates protein structural information into codon sequence generation. PISCO leverages a geometric deep learning architecture enabling the model to capture folding-aware translation dynamics. To enhance generalizability, we also introduce a species-agnostic variant that conditions on codon usage density, enabling optimization for previously uncharacterized or novel hosts.

We evaluate PISCO on a rigorously filtered benchmark and demonstrate that it outperforms state-of-the-art methods in codon recovery, distributional similarity, and local pattern alignment. Notably, PISCO achieves superior agreement with natural codon usage patterns, highlighting the importance of structure-aware inter-codon dependencies. Furthermore, we introduce a DNA quality assessment score to quantify the compatibility between coding sequences, protein structure, and the expression host. Finally, we validate PISCO through wet-lab experiments on multiple enzymes in carbon assimilation pathways. Our results show that structure-informed optimization leads to significantly improved protein solubility and functional expression in heterologous hosts. Collectively, PISCO provides a biologically informed toolset for precise genetic engineering, suggesting that codon optimization is fundamentally a structure-constrained and context-dependent process.

## 2 Results

### 2.1 Ablation analysis highlights the contributions of structural and host-specific features

Here, we empirically quantify the contribution of each feature—namely, species-name embeddings, secondary-structure-code embeddings, and ESM-derived protein-sequence embeddings—through an ablation study. To ensure comparability, we assess the pretrained models on the same hold out test set. As shown in Table 1, structural information and species-specific information emerge as the two main drivers of the improved recovery rate.

**Table 1:** An ablation study was conducted to quantify the marginal contribution of each individual feature.

| Method | Recovery↑ | MSE(CSI)↓ | MSE(CFD)↓ | DTW↓ |
| --- | --- | --- | --- | --- |
| PISCO-pretraining | **0.616** | **7.29E-3** | **21.4** | **0.158** |
| PISCO-pretraining w/o ESM <sup>1</sup> | 0.614 | 8.59E-3 | 22.0 | 0.177 |
| PISCO-pretraining w/o SSC <sup>2</sup> | 0.614 | 8.24E-3 | 23.5 | 0.164 |
| PISCO-pretraining w/o species name <sup>3</sup> | 0.568 | 11.1E-3 | 44.8 | 0.198 |
| PISCO-pretraining w/o protein structure <sup>4</sup> | 0.579 | 14.1E-3 | 26.2 | 0.226 |

<sup>1</sup>The residue level protein sequence embedding was not provided by the pretrained ESM2. It was replaced by an amino acid type embedding with same dimension.
<sup>2</sup>The Secondary Structure Code (SSC) was not provided.
<sup>3</sup>The species name embedding was not provided.
<sup>4</sup>Gaussian noise sampled from a distribution $N(0, 100)$ was introduced to the ESMFold predicted structures, and all associated features were updated accordingly. The model pretrained on this noise-augmented data was considered as not-incorporating explicit protein structural information.

### 2.2 PISCO outperforms SOTAs on a hold-out test set across multiple metrics

Building on this finding, we next evaluate whether explicitly modeling structural constraints can translate into stronger performance on a rigorously filtered hold-out benchmark spanning multiple enzyme families^1^.

As summarized in Table 2, all PISCO variants achieve higher codon recovery rates than state-of-the-art (SOTA) sequence-centric models^2^. Moreover, the codon usage distribution of PISCO-optimized sequences more closely resembles natural sequences, as indicated by lower mean squared errors in both the Codon Similarity Index (CSI) and Codon Frequency Distribution (CFD). Beyond single-codon accuracy, PISCO also better preserves local codon patterns, evidenced by reduced Dynamic Time Warping (DTW) distances relative to natural sequences.

**Table 2:**
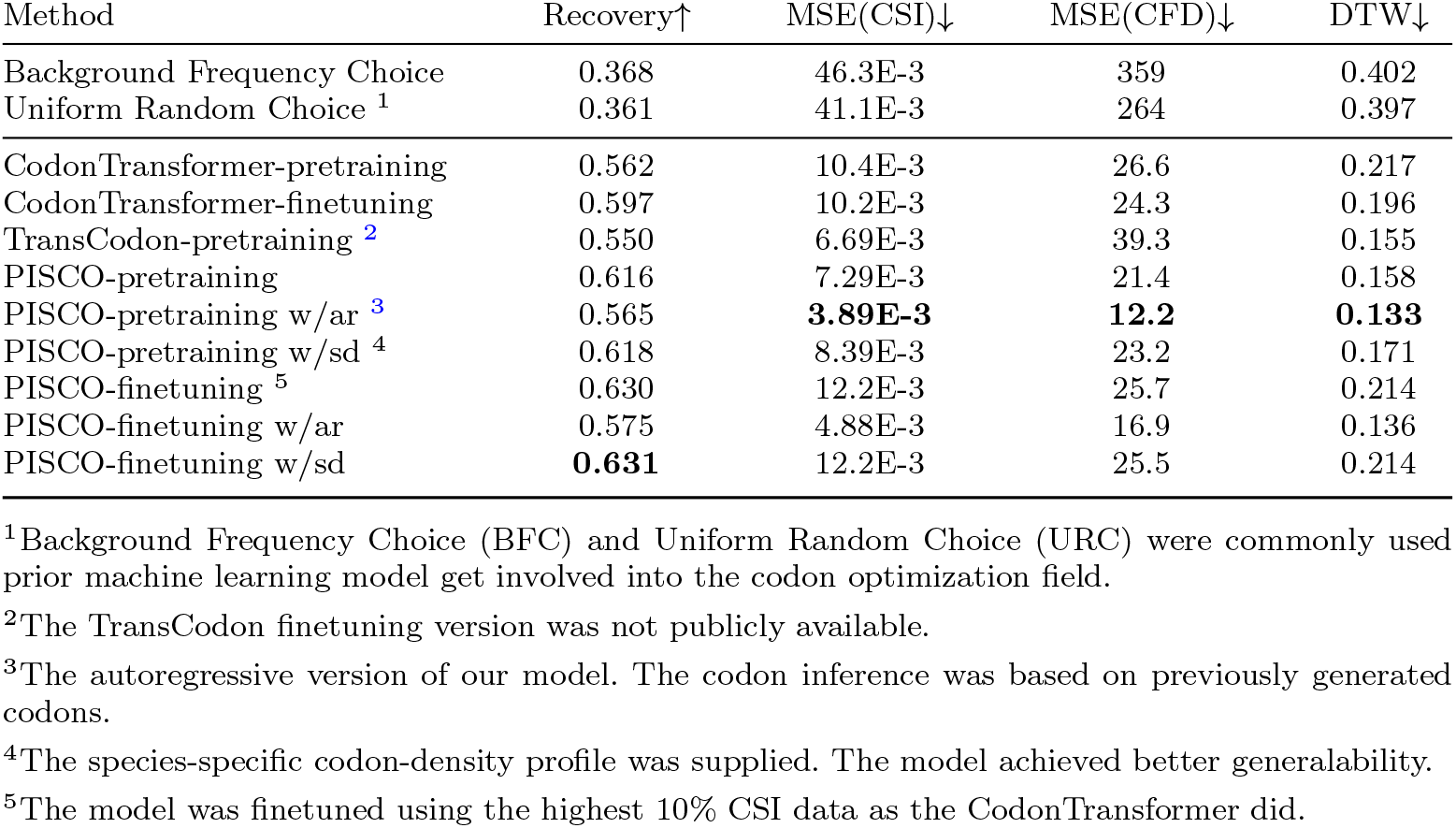
The codon recovery rate, Mean Square Error (MSE) of the Codon Similarity Index (CSI) and Codon Frequency Distribution (CFD) between the natural and optimized sequences were calculated. And the Dynamic Time Warping (DTW) between the natural and optimized sequences were calculated. The average of each evaluation metric across the 15 species in the test set were reported. The best value of each metric was bolded.

Two key observations emerge from these results. First, the autoregressive (AR) variant of PISCO, which conditions each codon prediction on preceding codons, achieves superior performance on distributional metrics (CSI, CFD, DTW) while showing a modest decrease in per-codon recovery. This suggests that leveraging contextual codon dependencies improves global similarity to natural sequences at the cost of a slight reduction in individual codon accuracy. Second, fine-tuning PISCO enhances codon recovery rates but may compromise distributional fidelity, highlighting a trade-off between single-codon accuracy and overall codon usage patterns.

### 2.3 PISCO captures both global and local codon usage patterns

Beyond per-codon recovery rates, the fidelity of codon optimization also depends on how well models reproduce global codon usage patterns and nucleotide composition constraints. To assess global patterns, we computed species-specific Codon Similarity Index (CSI), Codon Frequency Distribution (CFD), and GC content consistency for all models on the hold-out test set. As shown in Figures 2 and 3, most PISCO variants closely approximate the distributions observed in natural sequences. In particular, the autoregressive (AR) model achieves the highest degree of concordance across species in both codon usage and GC composition, indicating its ability to jointly capture codon bias and underlying nucleotide constraints. These results reinforce the observation from Table 2 that PISCO better captures host-specific codon preferences at the sequence level.

**Fig. 1:**
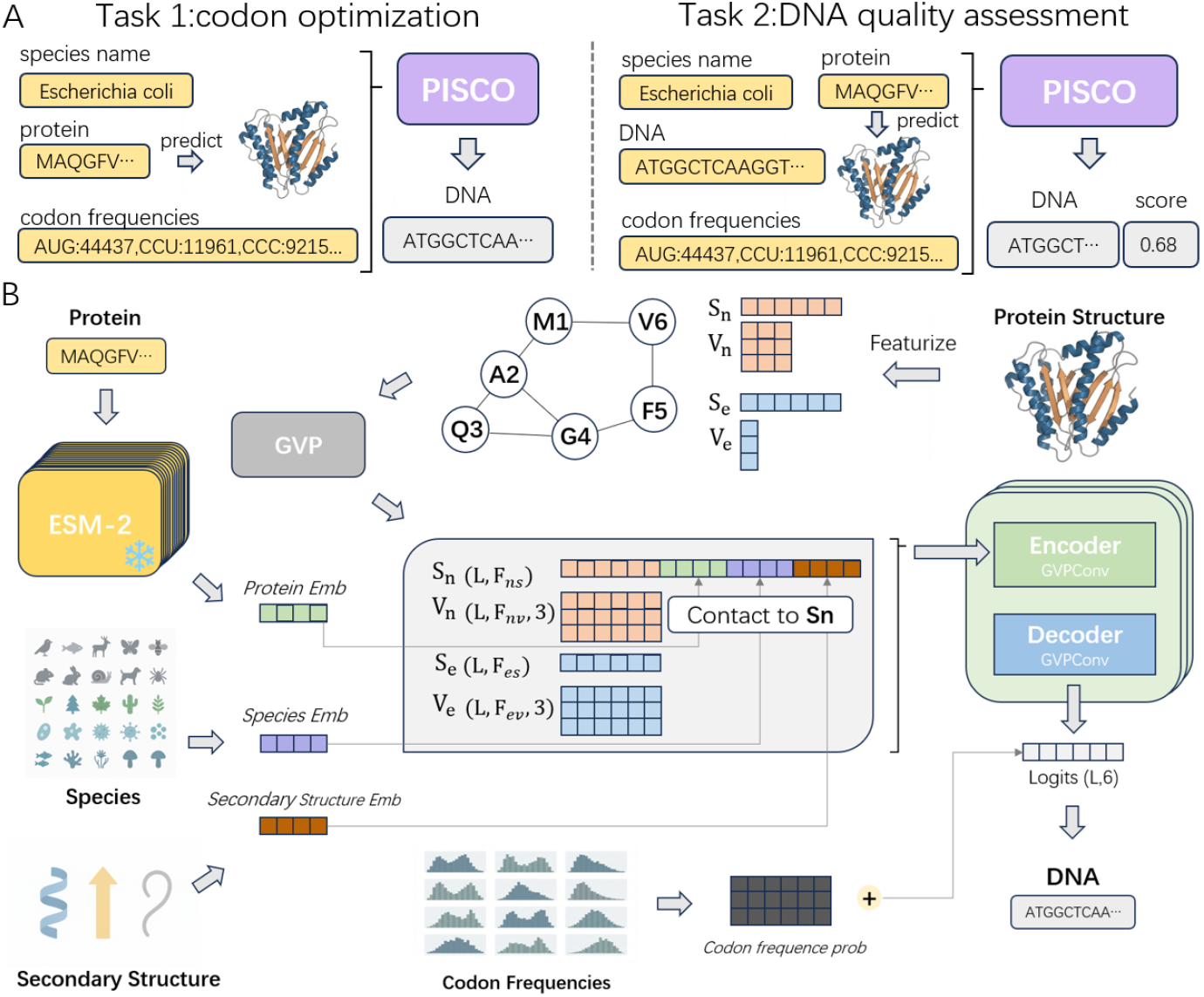
(A). Overview of PISCO tasks: 1.Codon optimization conditioned on host species, protein sequence and species-specific codon frequencies. 2. DNA quality assessment. Evaluate the compatibility among host species, heterologous DNA sequences and protein sequence. A lower score indicates greater mismatching and lower protein expression. In such cases, the use of our codon optimization model is strongly recommended. (B). Detailed model architecture. Inputs (protein sequence, inferred protein structure and host species) are encoded using the Geometric Vector Perceptron (GVP). The GVP processes inputs through parallel scalar and vector channels. The output is concatenated with a host-specific codon usage bias vector. Then used to predict the optimized DNA sequences (and quality scores).

**Fig. 2:**
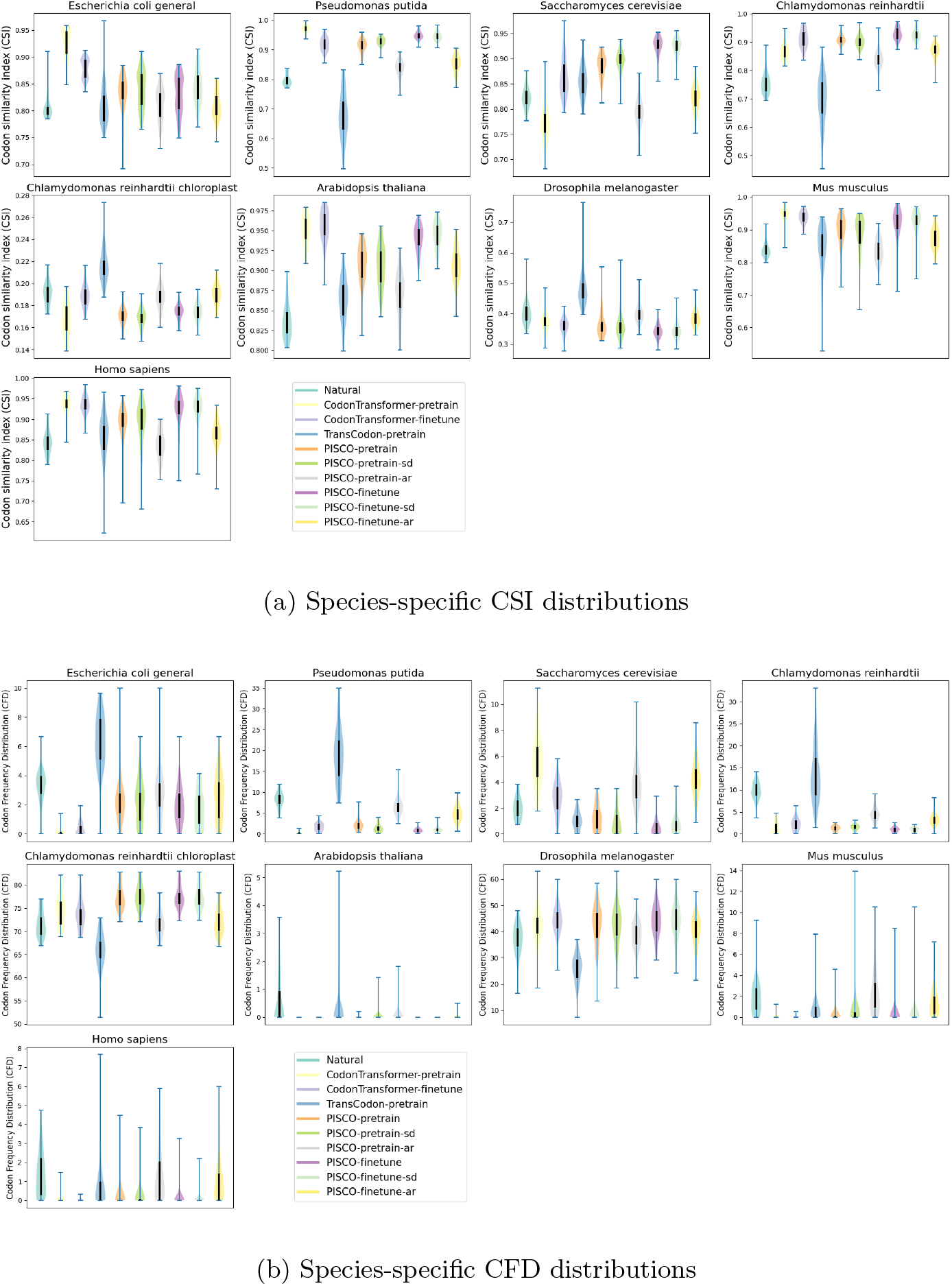
Species-specific distributions of global codon usage patterns across models. We utilized the nine species selected by CodonTransformer and generated violin plots to visualize the distributions of Codon Similarity Index (CSI) and Codon Frequency Distribution (CFD). Each subplot corresponds to a single species, where each violin represents a different model. The width of each violin reflects the probability density, and the thick solid line indicates the interquartile range of the model-specific distribution.

**Fig. 3:**
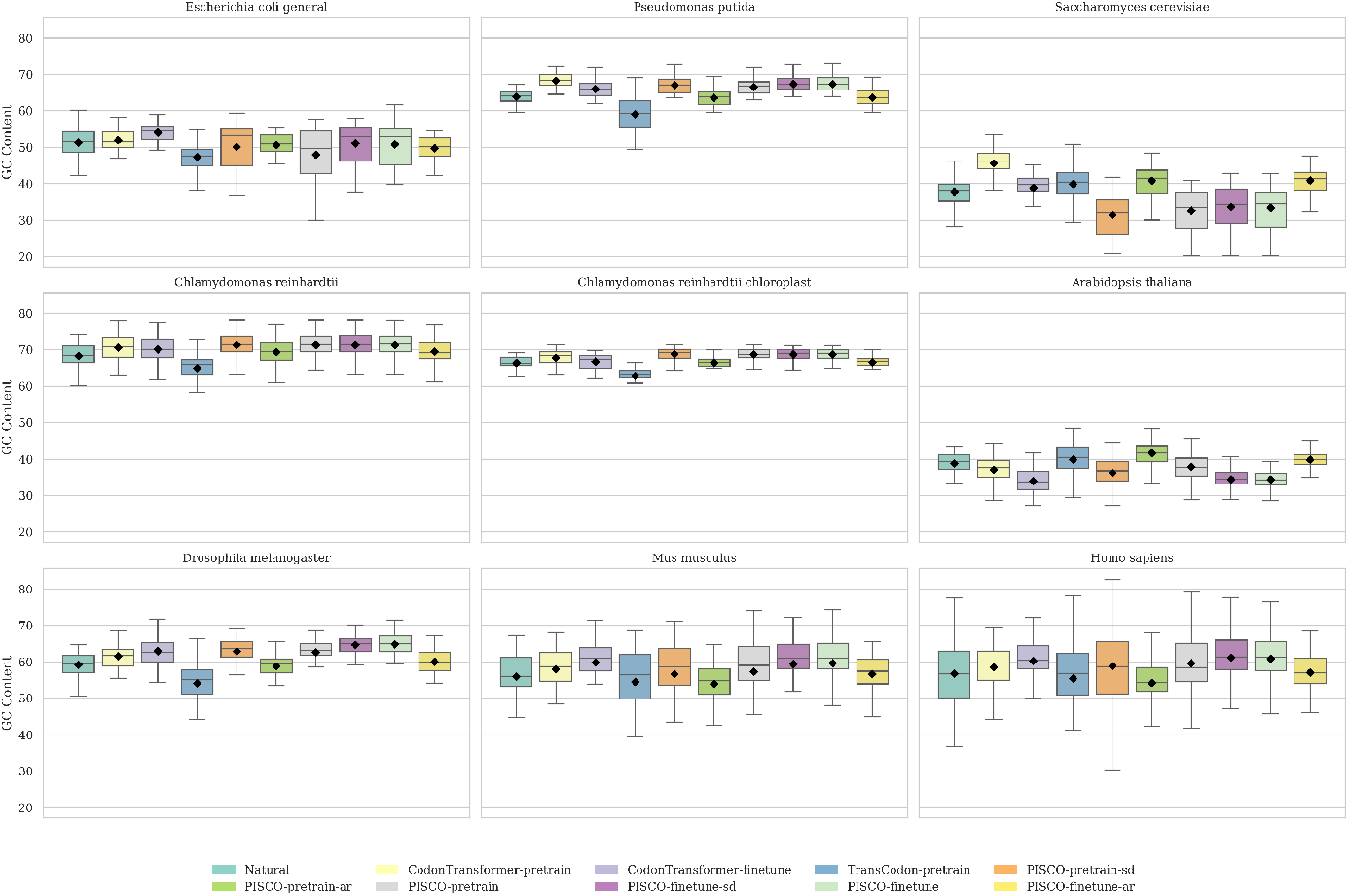
Species-specific distributions of GC content across models. We present boxplots to compare GC content distributions among different models across nine species. Each subplot corresponds to a single species, and each box represents a model. The box indicates the interquartile range, the central line denotes the median, and the diamond marker represents the mean GC content. This visualization highlights the ability of different models to preserve species-specific nucleotide composition biases.

While CSI, CFD, and GC content measure global distributional similarity, local codon arrangements are critical for co-translational folding. To evaluate local patterns, we computed Dynamic Time Warping (DTW) distances and visualized %MinMax curves using a sliding window of 18 codons. As illustrated in Figure 4, PISCO-AR exhibits the closest local codon patterns to natural sequences across multiple species, further highlighting the benefits of context-aware codon prediction.

**Fig. 4:**
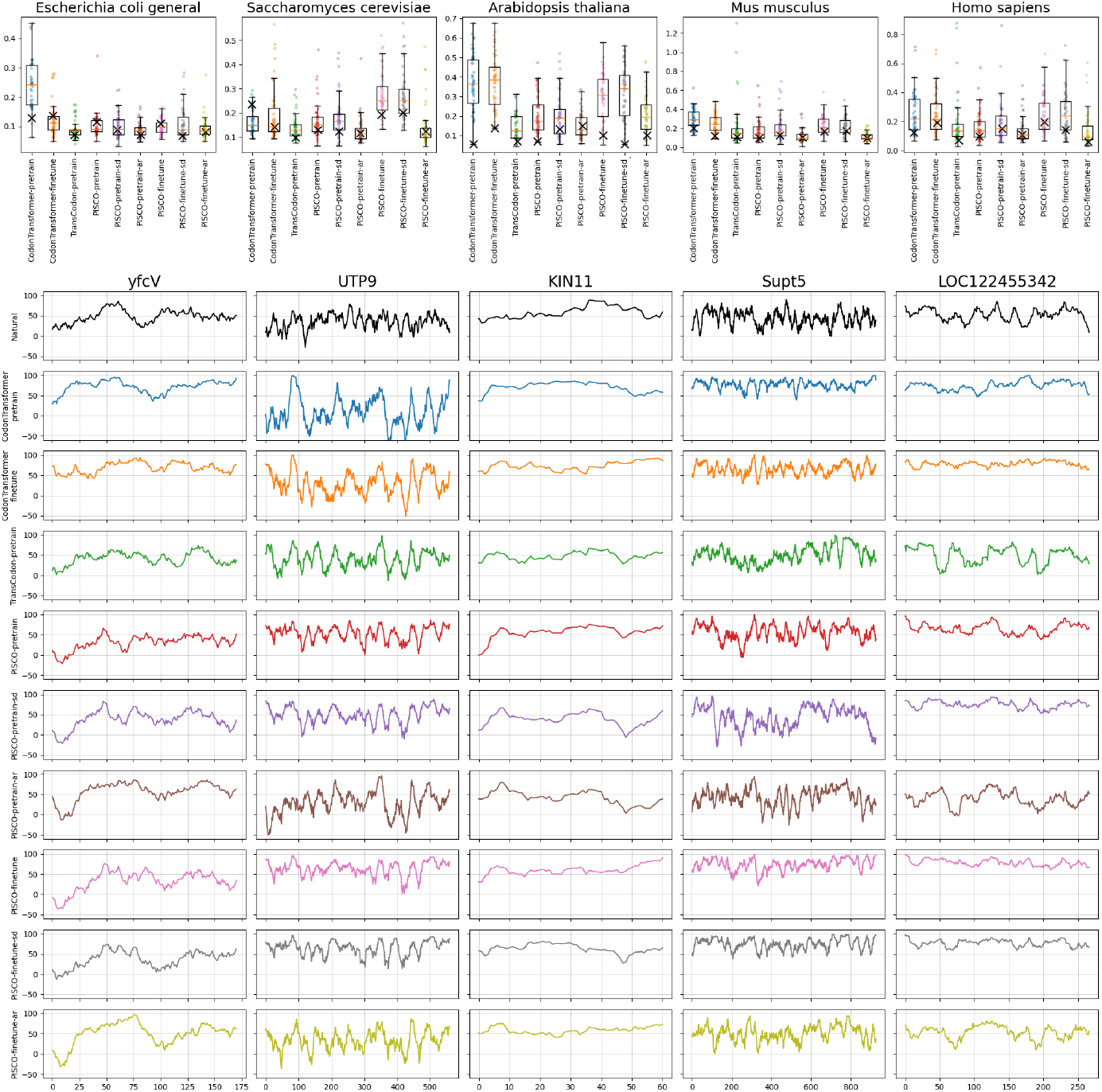
We use the same 5 species selected by the CodonTransformer and draw the boxplot of DTW of optimized codon sequences to natural sequences. Each subplot is for one species. And each box is representing model. We randomly select one sample from each species, marked as ‘x’ in boxplot. And visualize the %*MinMax* of the selected gene. The gene name is shown on the top of each column.

Taken together, these analyses demonstrate that PISCO effectively captures not only global codon usage patterns and nucleotide composition (GC content), but also local sequence motifs, underscoring the importance of integrating structural and host-specific information.

### 2.4 PISCO DNA assessment score can successfully discriminate heterologous DNA sequences

To evaluate the performance of our model in DNA quality assessment, we select the industrially-common yeast Komagataella pastoris (formerly Pichia pastoris). 15 genes are randomly chosen from K. pastoris, covering different chromosomes and protein functions. For each gene, a homologous protein from another species is randomly selected, with the constraint that the sequence identity falls between 50% and 90%. The lower bound ensures functional similarity, while the upper limit excludes proteins from overly related species where codon optimization is less necessary for heterologous expression. As K. pastoris is not included in the training data set, species-specific inference was performed using our model with the sd variant. The DNA + protein sequence + protein structure + species codon density matching scores is predicted. A higher score indicates a better match and a high protein expression level. As shown in the Fig. 5, the retraining model accurately prioritize 12/15 of the genes originating from K. pastoris, and finetuning model accurately prioritize 13/15 of the genes. Finetuning prioritize PRS26A, which is marginally better than pretraining model.

**Fig. 5:**
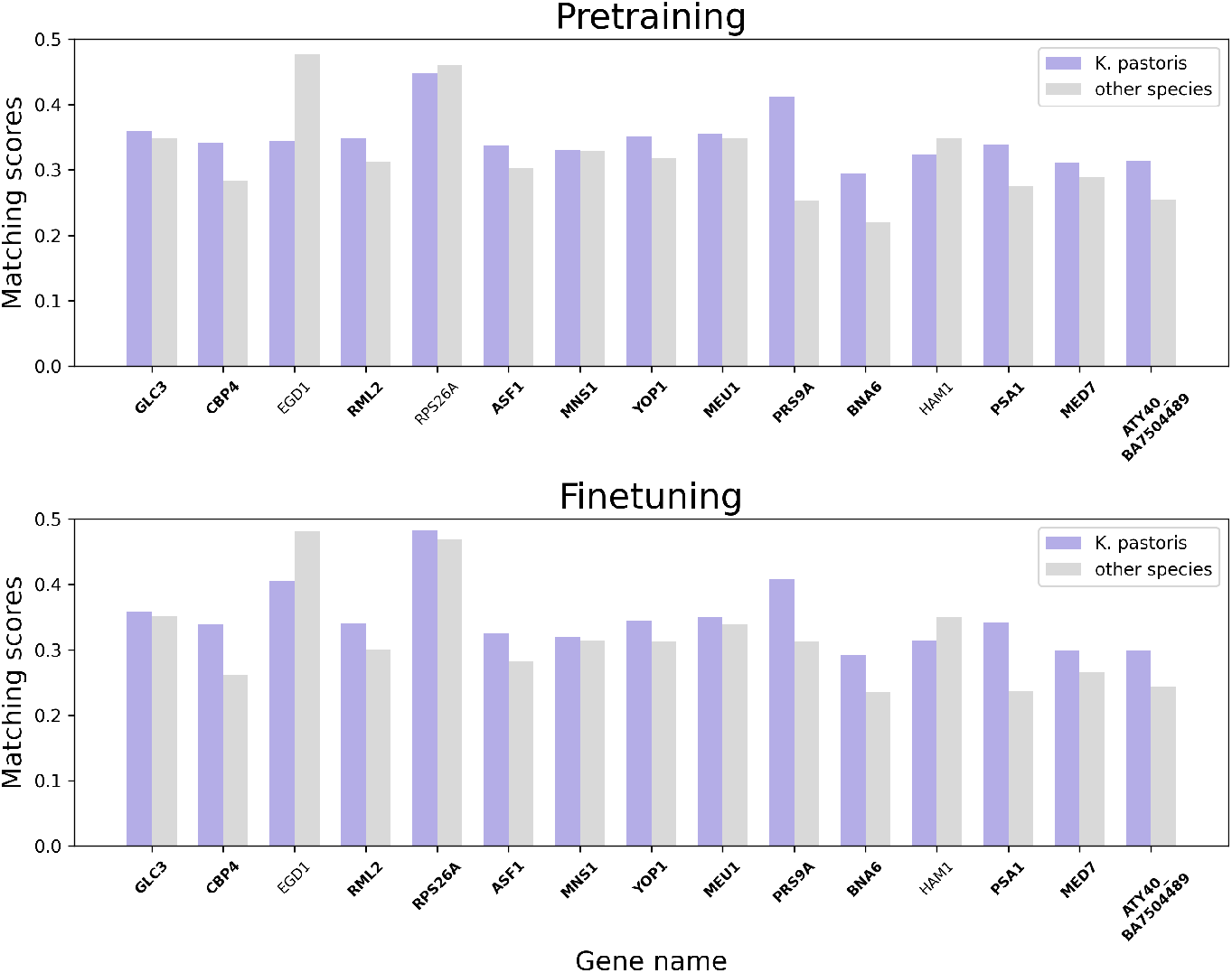
We predicted the DNA + protein sequence + protein structure + species codon density for each selected gene. The DNA sequences originated from K. pastoris should have higher predicted scores than their heterologous counterparts.. The genes our model assign higher score to the K. pastoris sequence were in bold.

### 2.5 PISCO optimized and prioritized codon sequences have high heterologous expression

We select three key enzymes—Ribulose-1,5-bisphosphate carboxylase/oxygenase (RuBisCO), Phosphoribulokinase (PRK), and Formate Dehydrogenase (FDH)—as optimization targets, as they constitute the core modules for carbon assimilation and energy generation in synthetic autotrophic yeast[19, 20]. Specifically, PRK and RuBisCO facilitate the heterologous Calvin-Benson-Bassham (CBB) cycle to enhance carbon yield: PRK converts D-ribulose-5-phosphate into Ribulose-1,5-bisphosphate (RuBP) via ATP-dependent phosphorylation, followed by RuBisCO fixing CO2 into RuBP. To sustain this energy-intensive process, FDH catalyzes the oxidation of formate to CO2 to generate the NADH required for biomass formation while preventing toxic formate accumulation. Homologous sequences of RuBisCO and PRK from various species are retrieved from the UniProtKB database[19].

Codon optimization is performed using PISCO finetuning w/sd model. The optimized sequences are ranked accordingly. The top 5 RuBisCO and top 13 PRK codon-optimized sequences are selected for wet-lab experimental validation. A reference set of RuBisCO (UniProt: Q60028) and PRK (UniProt: P09559), previously reported to be effective for the engineering of autotrophic K. pastoris, was selected as the control group to evaluate the expression enhancement provided by PISCO-generated sequences[19]. RuBisCO and PRK variants are designated by numerical identifiers due to proprietary considerations. As shown in Figures 6, Western blot analysis confirmed abundant and soluble expression levels for the majority of the selected sequences. Clear, distinct bands were observed at the expected molecular weights (52.5-52.9 kDa for RuBisCO, 24.5-47.2 kDa for PRK and 40.4-41.1 kDa for FDH). Relative expression levels were determined via densitometry using ImageJ[21], calculated as the ratio of the target band density to the control sample. For RuBisCO, the codon-optimized variant RuBisCO4 exhibited superior performance in both soluble and total protein expression compared to the Q60028 control. Among the 13 PRK variants, 12 exhibited higher soluble expression levels than the P09559 control, with the sole exception of PRK3. For FDH, five out of seven optimized variants exhibited substantial soluble expression levels, with the top-performing candidate A0A395MPN4 showing 2.3-fold improvement in total protein yield compared to the unoptimized control sequence. More details can be found in section 4.2.

**Fig. 6:**
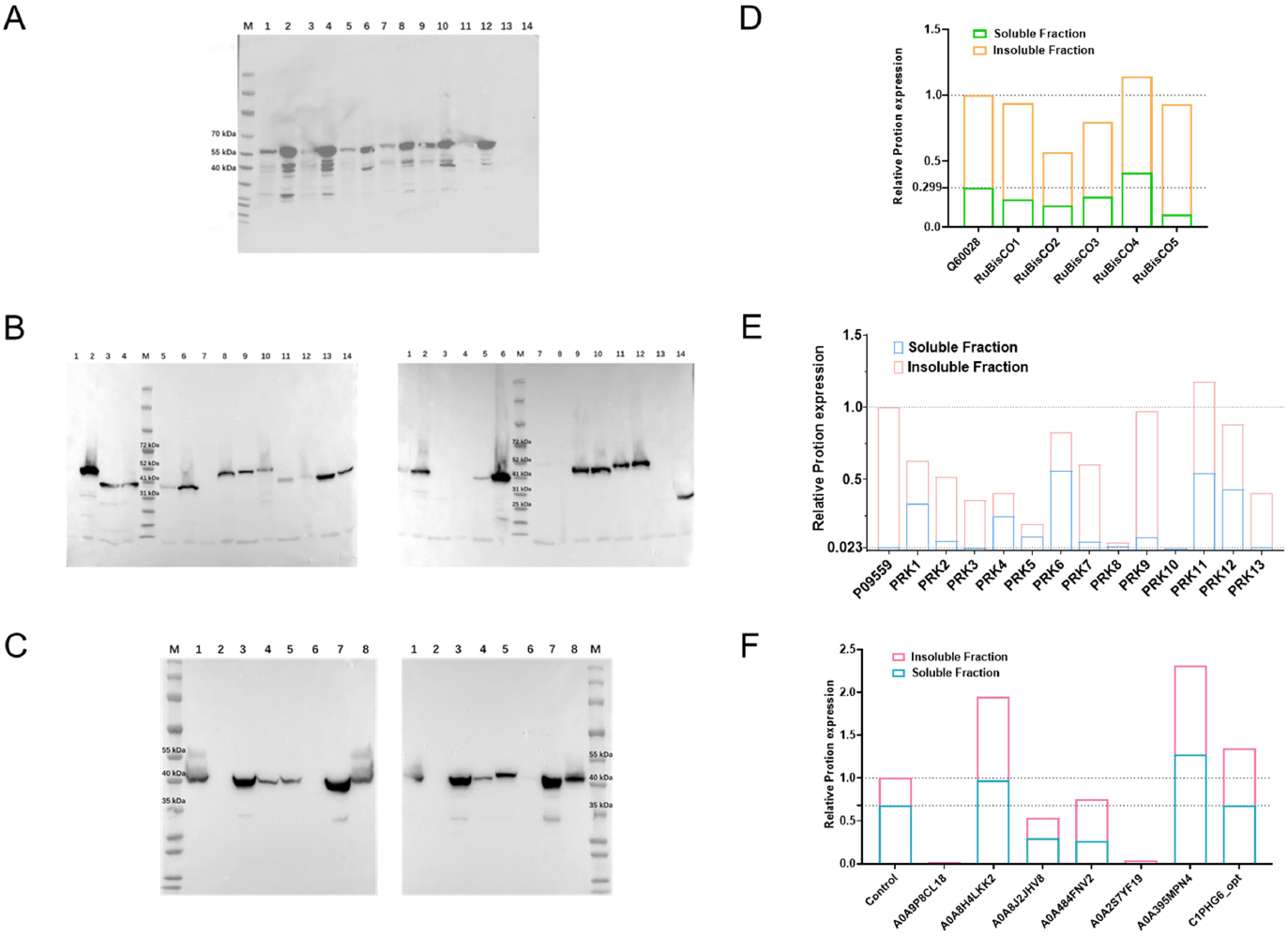
Heterologous expression and quantitative analysis of PISCO-optimized sequences in K. pastoris. (A) Western blot analysis of five RuBisCO variants prioritized by PISCO-finetuning. Molecular weight markers (M) indicate RuBisCO at approximately 53 kDa. The loading order corresponds directly to panel D. For each variant, two adjacent lanes are presented: the left lane contains the soluble protein fraction (supernatant), while the right lane contains the insoluble fraction (cell debris). (B) Western blot analysis of 13 PRK variants, with bands appearing within the expected molecular weight range of 24.5–47.2 kDa. The loading order corresponds directly to panel E. For each variant, adjacent lanes display the soluble (left) and insoluble (right) fractions, respectively. (C) Western blot analysis of eight FDH variants. Molecular weight markers indicate FDH at approximately 41 kDa. For each variant, lanes on the left represent the soluble fraction (supernatant), whereas lanes on the right represent the insoluble fraction (cell debris). (D–F) Relative protein expression levels of RuBisCO (D), PRK (E), and FDH (F) quantified by densitometric analysis of Western blot bands using ImageJ. Stacked bars represent total protein expression, with the lower portion indicating the soluble fraction and the upper portion indicating the insoluble fraction. All data were normalized to the respective control values.

To assess the correlation between PISCO scores and actual protein expression, we performed correlation analysis for PRK variants using the experimentally validated dataset. As shown in Figures 7, scatter plots illustrate the relationship between PISCO scores and both soluble protein expression (Fig. 7A) and total protein expression (Fig. 7B), with Pearson correlation coefficients calculated to quantify these associations. The moderate correlation observed can be attributed to two primary factors. First, the limited sample size (n=13 for PRK) constrains statistical power. Second, as we selected only high-scoring candidates for experimental validation, the restricted score range within this top-tier subset inherently attenuates the observable correlation between minor score variations and expression levels.

**Fig. 7:**
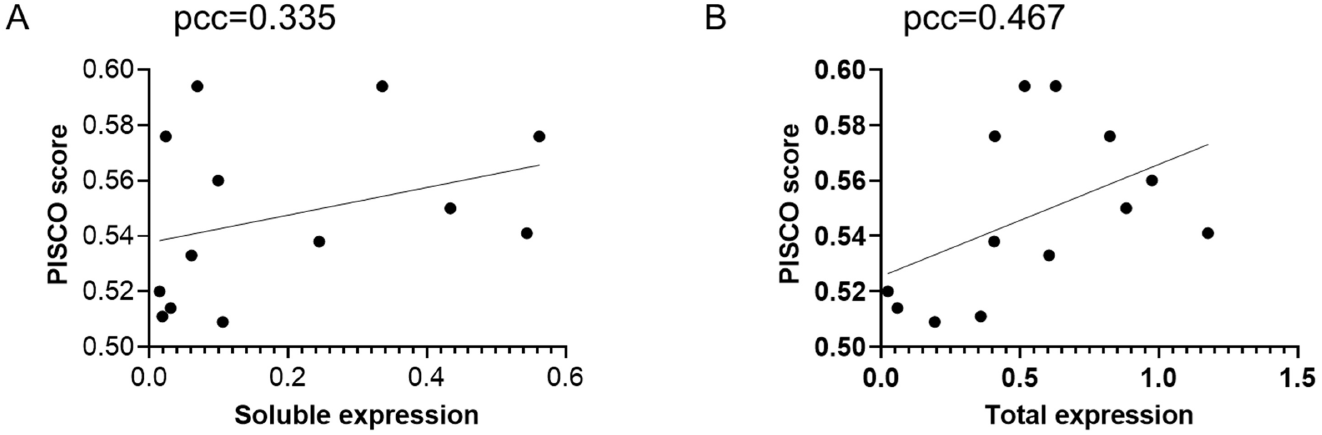
Correlation analysis between PISCO score and protein expression levels of PRK. (A) Scatter plot showing the relationship between soluble protein expression level (x-axis) and PISCO score (y-axis). Pearson r = 0.335, p = 0.263. (B) Correlation between total protein expression level (combined soluble and insoluble fractions, x-axis) and PISCO score (y-axis). Pearson r = 0.467, p = 0.108.

We evaluated the correlation between different codon optimization metrics and experimental protein expression outcomes, including soluble and total expression (Table 3).The Natural Codon Similarity Index (CSI) is computed based on codon usage frequencies derived from *K. pastoris*, serving as a frequency-based baseline for host adaptation. In contrast, we compare two variants of our method: PISCO-ar, an autoregressive formulation, and PISCO-sd, a synchronous decoding variant, to explore different modeling strategies for codon optimization.As shown in Table 3, PISCO-sd achieves the highest correlation with both soluble (*r* = 0.335) and total expression (*r* = 0.467). CSI shows weak or negligible correlation, particularly for total expression (*r* = −0.053), while PISCO-ar exhibits limited correlation overall. Although the correlations are not statistically significant (*p >* 0.05), likely due to the limited sample size, the results suggest that PISCO-sd provides a more informative signal for predicting expression outcomes compared to both frequency-based and autoregressive approaches.

**Table 3:** Correlation between different scoring metrics and protein expression outcomes.

| Metric | Soluble |  | Total |  |
| --- | --- | --- | --- | --- |
| | $r$ | $p$ | $r$ | $p$ |
| CSI | 0.267 | 0.378 | -0.053 | 0.865 |
| PISCO-ar | 0.032 | 0.916 | 0.273 | 0.367 |
| PISCO-sd | **0.335** | 0.263 | **0.467** | 0.108 |

## 3 Discussion

Our results suggest that PISCO is not merely a codon substitution tool, but a model that learns a biologically meaningful “codon language” shaped jointly by host-specific preferences, protein structural constraints, and codon-context dependencies. This perspective helps explain why PISCO can simultaneously improve codon recovery, preserve natural codon usage patterns, and identify sequences with higher expression potential.

Ablation analysis indicates that species identity and protein structural information are the dominant contributors to model performance. Removing either of these inputs causes a clear degradation in recovery, CSI, CFD, and DTW, whereas removing residue-level ESM embeddings or secondary structure codes leads to only modest performance drops. These results imply that effective codon optimization depends primarily on two sources of information: the host-specific codon preference landscape and the structural context of the encoded protein. In other words, codon choice is not determined by frequency statistics alone, but is also constrained by protein-level organization that may influence translation and folding.

Consistent with this view, PISCO outperforms existing sequence-centric baselines on the hold-out test set across both recovery and distributional metrics. Beyond recovering individual codons, the model better reproduces species-specific CSI and CFD distributions and achieves lower DTW distances, showing that it captures both global codon usage bias and local codon arrangement patterns. The autoregressive variant further strengthens this effect, especially on CSI, CFD, and DTW, suggesting that conditioning each prediction on previously generated codons helps the model learn contextual codon dependencies that better approximate natural sequences. At the same time, the slight reduction in per-codon recovery indicates a trade-off between exact codon matching and distributional fidelity.

The species-specific codon-density profile variant of PISCO, denoted as PISCO-sd, plays a distinct role in this framework. By supplying external species-level codon density information, PISCO-sd is able to inject codon preference signals for species that were not seen during training, thereby improving generalization to unseen hosts. This design is particularly important because it allows the model to adapt to host-specific codon bias even when the target species is absent from the training distribution. In our experiments, PISCO-sd was also the version used for downstream scoring and wet-lab prioritization, indicating that it is especially suitable for ranking candidate sequences by their expected expression performance.

The DNA assessment experiments further support the generalizability of the learned scoring function. When applied to *Komagataella pastoris*, which was excluded from training, PISCO correctly assigned higher scores to native DNA sequences than to homologous heterologous counterparts in most cases. This suggests that the score reflects more than simple training-set memorization: it captures sequence quality in a way that transfers to unseen species. Such behavior is desirable for practical codon optimization, where the ability to evaluate sequences in new hosts is often more valuable than merely reproducing patterns observed in the training data.

Most importantly, wet-lab validation confirms that these computational scores are biologically meaningful. PISCO-sd-guided prioritization of RuBisCO, PRK, and FDH variants led to strong heterologous expression in *K. pastoris*, with many optimized sequences outperforming established control constructs. The majority of selected variants showed abundant soluble expression, and the best FDH candidate achieved a substantial increase in total protein yield. These results demonstrate that PISCO is not only capable of generating codon-optimized sequences that resemble natural ones, but also capable of prioritizing candidates that translate into improved experimental outcomes. Correlation analysis between PISCO scores and measured expression levels further supports this conclusion: although the correlations are moderate and not statistically significant due to the limited sample size and the restricted score range of the selected candidates, PISCO-sd still shows the strongest association with both soluble and total expression among the compared metrics.

Taken together, these findings support a unified interpretation of PISCO as a host- and structure-aware codon optimization framework. The model learns to balance natural codon usage patterns, protein structural constraints, and host-specific preferences, while the species-density-based formulation extends its applicability to unseen hosts. The resulting scoring function is not only biologically interpretable, but also practically useful for ranking and selecting heterologous coding sequences with improved expression potential.

## 4 Methods

### 4.1 Data

We used the dataset curated by CodonTransfromer, which contains 1,001,197 DNA sequences derived from 164 species.[15] Consistent with the original study’s protocol, we adopted the same splitting method for pre-training and fine-tuning subsets. Amino acid sequences, translated from the corresponding DNA sequences, are used as input for ESMFold[22] to predict their three-dimensional structures. To mitigate ESMFold prediction failures and prevent out-of-memory errors, sequences yielding translations exceeding 1,000 amino acids were excluded. Protein sequences shorter than 15 amino acids were also excluded to ensure that the corresponding peptides possess a stable, well-defined secondary structure.[23] Following the same evaluation protocol used by CodonTransformer, model performances were evaluated using the same 15 selected species. 50 samples from each species were selected, except the *Chlamydomonas reinhardtii chloroplast*, which only had 28 samples after applying filtration. So, in total 728 (=14 * 50 + 28) samples were selected as the holdout test. To prevent information leakage between training and test sets, we excluded all samples whose protein similarity to any sequences in the holdout test that exceeded 40%. To minimize the impact of the sequence excluding, we choose the 728 test sequences with minimum sequence excluding needed. Protein sequence similarity is calculated using mmseqs2.[24] After applying the sequence length filter and the protein similarity filter, 883,595 pre-training and 41,012 fine-tuning samples were left. 1,000 samples were randomly selected from the finetuning dataset as the validation for both pretraining and finetuning.

### 4.2 K. pastoris heterologous expression experiment

The DNA sequences of the selected RuBisCO, PRK and FDH variants were synthesized and cloned into the standard expression vector *pPICZα* under the transcriptional control of the methanol-inducible AOX1 promoter. All plasmids were synthesized and verified by Tsingke Biotechnology Co., Ltd. (Beijing, China). The recombinant plasmids were linearized via digestion with SacI and transformed into K. pastoris strain X-33 using electroporation. Transformants were screened on selective YPD agar plates containing Zeocin (100 *µ*g/mL; Solarbio, China). For protein expression, positive clones were initially cultured in YPD medium at 30°C with shaking at 220 rpm until the *OD*_600_ reached 4-6. The cells were then inoculated at an 1% (v/v) ratio into 50 mL of BMMY (Buffered Methanol-complex Medium) to initiate induction. Methanol was supplemented to a final concentration of 2% (v/v). The compositions of YPD and BMMY media were prepared according to standard protocols[25].

After 48 hours of induction, cell pellets were harvested via centrifugation. Total intracellular protein was extracted via freezing in liquid nitrogen followed by mechanical grinding. Microscopic examination of the ground samples confirmed that the vast majority of cells were fragmented, ensuring efficient protein recovery. Protein expression was analyzed via Western blot. Specifically, 20 *µ*L protein sample was mixed with 5 *µ*L loading buffer. Samples were resolved using 8-16% precast SDS-PAGE gels (GenScript, China) and subsequently transferred to PVDF membranes (Invitrogen). The membranes were blocked with 5% non-fat milk for 1 hour and incubated with HRP-conjugated anti-His-tag antibody (for RuBisCO and FDH) or HRP-conjugated anti-Myc-tag antibody (for PRK) for 1 hour. Protein bands were visualized using enhanced chemiluminescence (ECL) to assess the expression and solubility profiles.

### 4.3 Model

We propose **PISCO** (**P**rotein-**I**nformed **S**pecies-aware **C**odon **O**ptimizer), a structure-aware neural network for codon prediction conditioned on protein sequence, three-dimensional structure, and species-specific codon usage bias. Given a protein of length *L*, the model predicts, for each residue position(predicted by ESMFold[22]), a probability distribution over synonymous codons in a fully parallel manner.

### 4.4 Protein Graph Representation

A protein is represented as a residue-level graph *G* = (*V, E*), where each node corresponds to an amino acid residue and edges encode geometric relations derived from the 3D structure. Each node *i* is associated with scalar–vector features 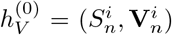,and each edge (*i, j*) is associated with 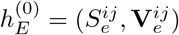.

#### Initial Feature Projection

The original node and edge features consist of low-dimensional scalar and vector components. Specifically, the node scalar and vector features satisfy 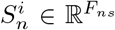 and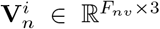, while the edge scalar and vector features satisfy 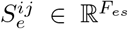 and 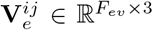. In our implementation, the input dimensions are *F*_*ns*_ = 6, *F*_*nv*_ = 3, *F*_*es*_ = 32, and *F*_*ev*_ = 1(as described in [26]). These features are projected into a higher-dimensional latent space using Geometric Vector Perceptrons (GVPs), resulting in node features with *F*_*ns*_ = 64 and *F*_*nv*_ = 16, and edge features with *F*_*es*_ = 32 and *F*_*ev*_ = 6. Layer normalization is applied to the projected representations to stabilize training and ensure consistent scaling across scalar and vector channels.

### 4.5 Encoder–Decoder Architecture

Both **PISCO** and its autoregressive variant **PISCO-ar** adopt a GVP-based encoder– decoder architecture operating on residue-level protein structure graphs. The encoder consists of stacked GVP convolution layers that propagate geometric messages across the residue graph, producing structure-aware residue embeddings.

Formally, given residue features *h*_*V*_ and geometric edge features *h*_*E*_, the encoder updates node states as:

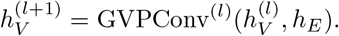

#### PISCO (Synchronous Decoding)

PISCO decodes all residues in parallel, without modeling inter-codon dependencies. The decoder mirrors the encoder while conditioning on encoder outputs:

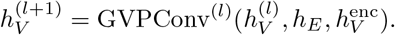

A linear head then predicts codon probabilities synchronously for all positions:

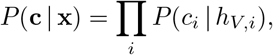

This enables efficient parallel inference with a single forward pass.

#### PISCO-ar (Autoregressive Decoding)

PISCO-ar introduces codon-aware dependencies via causal message passing. During training, ground-truth codons **c** are embedded and attached to forward edges (source index *<* destination index), forming codon-augmented edge features and enabling parallel teacher forcing. At inference time, decoding becomes autoregressive:

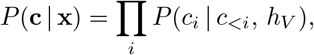

where previously predicted codons *c*_*<i*_ are embedded and propagated along valid forward edges, while unresolved nodes remain masked. This graph-based causal masking preserves biological ordering and modulates the structural context with codon history.

### 4.6 Feature Fusion

All scalar features are concatenated to form the final node scalar representation

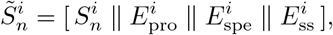

while the vector features remain unchanged. The fused node representation is then given by 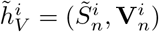, which is used as the input to subsequent GVP-based encoder and decoder layers.

#### Amino Acid Embedding

Residue-level sequence information is incorporated using contextual embeddings from a pretrained ESM2[22] model. Since the original ESM2 embeddings are high-dimensional and may dominate other feature channels when directly concatenated,we apply a linear projection to reduce their dimensionality and balance their contribution with structural and species-related features. The projection dimension *d*_pro_ is selected as 128 based on extensive hyperparameter tuning. Let 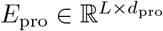denote the resulting per-residue protein sequence embeddings.

#### Species Embedding

Each protein is associated with a species identifier *s*. A trainable species embedding 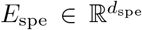 is learned and broadcast to all residues. During training, the species identifier is randomly masked with a small probability to improve robustness to missing or uncertain species annotations.

#### Secondary Structure Embedding

If available, residue-level secondary structure annotations (coil, helix, and sheet) are embedded using a trainable embedding layer, yielding representations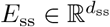.

### 4.7 Codon Prediction and Constraints

#### Prediction Head

Although there are sixty-four sense codons in the standard genetic code, the number of synonymous codons varies across amino acids and is upper-bounded by six. We therefore formulate codon prediction as a grouped classification problem conditioned on the amino acid identity. Specifically, a final GVP layer maps each residue representation to a 6-dimensional logit vector logits_*i*_ ∈ℝ^6^, where each dimension corresponds to one of the possible synonymous codon slots for the given amino acid. This design enables a unified output space across all residues while avoiding redundant parameters associated with a full codon vocabulary.

#### Amino Acid–Codon Masking

To enforce biological validity, logits corresponding to codons incompatible with the residue’s amino acid identity are masked with −∞ prior to normalization, ensuring that only synonymous codons receive non-zero probability mass.

#### Species-Specific Codon Usage Prior

Optionally, PISCO incorporates species-specific codon usage bias by adding a prior distribution *P*_spe_(codon | *a, s*) in log-space to the predicted logits, i.e.,

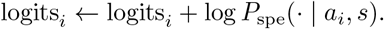

The model is trained using codon usage statistics derived from 164 species. To support codon optimization and DNA quality assessment for arbitrary species beyond the training set, PISCO is designed to dynamically load external codon usage statistics at inference time, enabling flexible adaptation to unseen species without modifying model parameters.

### 4.8 Training

Training was performed in two stages: pretraining from scratch followed by full fine-tuning on the highest 10% CSI data. During pretraining, PISCO was trained on protein structure graphs for codon prediction, taking residue-level structural graphs as input and predicting synonymous codons using a cross-entropy objective. The model was optimized using Adam with dynamic graph batching, where each batch was constrained to a maximum of 20,000 nodes. Pretraining was conducted for 23 epochs on a single NVIDIA A100 GPU with 80 GB memory, requiring approximately 85 hours in total. For finetuning, mixed-precision training was employed on the same hardware. Models were optimized using AdamW with an initial learning rate of 1×10^*−*4^ and a OneCycle learning rate schedule with a warm-up fraction of 0.1, updated at every training step. Finetuning was performed for 5 epochs using dynamic graph batching with a maximum of 20,000 nodes per batch.

### 4.9 Evaluation metrics

The following several metrics are used to evaluate the optimized DNA sequences.

#### 4.9.1 Codon recovery rate

Codon recovery rate quantifies the percentage of codons that have been perfectly reconstructed. For a given codon sequence A comprising L codons, A=*{codon*_*i*_, *i* = 1, 2, …, *L}*, and its corresponding optimized codon sequences 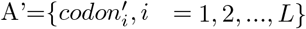.

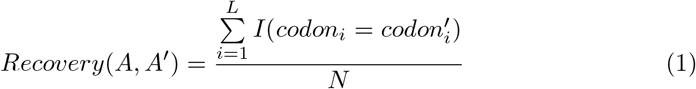

where *I* is the indicator function. *I* (*True*) = 1 and *I*(*False*) = 0.

#### 4.9.2 Codon similarity index (CSI)

The CSI quantifies the degree to which the codon usage of a sequence conforms to the genome-wide codon distribution of the reference species. CSI is a refinement of the commonly used metric codon adaptation index (CAI).[27] The CAI is predicted on a restricted set of highly expressed genes. To avoid bias caused by arbitrary selection, CSI[28] uses the genome-wide codon frequencies.

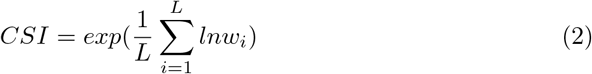

where the relative adaptiveness of a codon 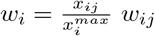 is calculated as the ratio of its frequency *x*_*ij*_ to the most frequent synonymous codon 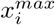.

#### 4.9.3 Codon frequency distribution (CFD)

CFD measures the usage of rare codons.

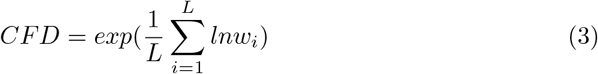

where *w*_*i*_ = 1 if the codon is rare. 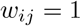 if 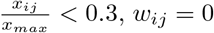 otherwise.

#### 4.9.4 Mean Square Error

To evaluate the performance of various codon optimization models with respect to CSI and CFD metrics, we employed the Mean Square Error (MSE) between natural and optimized codon sequences. The MSE is selected for two primary reasons. First, unlike the Kullback–Leibler (KL) divergence used by TransCodon [18], which quantifies similarity between distributions, MSE provides a direct, pairwise comparison. Second, compared to the Mean Absolute Error (MAE), MSE is more sensitive to outliers. For each species, the MSE was computed as the average squared error across all samples for either CSI or CFD.

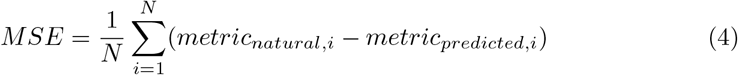

where, *metric*_*natural,i*_ and *metric*_*predicted,i*_ are the CSI or CFD values of the *ith* natural sequence and the model optimized sequence respectively.

#### 4.9.5 Dynamic time warping (DTW)

%*MinMax* evaluates the balance between high and low frequency codons within a sliding window along the sequence. The window size *w* is commonly used as 18, i.e. *w* = 18. The 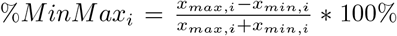. And the overall %*MinMax* of a gene is represented as an array.

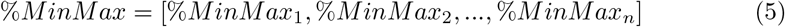

where n is the number of the windows of the given gene. The DTW evaluates the similarity between two temporal sequences. Here, DTW is used to measure the similarity between %*MinMax* array of the native codon sequence and the generated codon sequence, denoted as X and Y respectively.

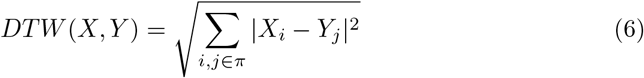

where *π* is the optimal alignment path of the X and Y. *X*_*i*_ and *Y*_*j*_ are the matched points in the aligned sequence. DTW calculated the Euclidean distance between those aligned points.

## 6 Code availability

The PISCO model and codes used to train and evaluate models are available at the GitHub repository: https://github.com/MoleculeMindOpenSource/PISCO.

## 7 Author contribution

D.Z. conceived the study. D.Z and W.T. constructed the model. W.J. implemented the code. W.J. and W.T. analyzed the model results. H.L. and X.J. performed the wet-lab experiment with M.L.’s help. All authors drafted the manuscript and approved the final draft.

## Footnotes

1 The benchmark is based on the dataset curated by CodonTransformer [15], with additional filtering by protein sequence identity (see Methods, Section 4.1).

2 Notably, we employed publicly released weights for CodonTransformer and TransCodon models, which may potentially exposed to the test samples. PISCO has no such data leakage, further validating its performance gains.

